# A temperature-dependent translational switch controls biofilm development in *Vibrio cholerae*

**DOI:** 10.1101/2020.05.07.082040

**Authors:** Teresa del Peso Santos, Jonathon Blake, Brandon Sit, Alyson R. Warr, Vladimir Benes, Matthew K. Waldor, Felipe Cava

## Abstract

The gastrointestinal pathogen *Vibrio cholerae* frequently forms biofilms during its life cycle. Biofilm formation is vital for protection against environmental stresses and is thought to facilitate intestinal colonization. Adaptation to temperature is crucial for *V. cholerae* survival, as the pathogen is exposed to seasonal temperature variations in the aquatic environment, and temperature fluctuations during host-environment transitions. Here, we show that *V. cholerae* strains naturally lacking the master biofilm transcriptional regulator HapR are unable to develop colony rugosity at low temperatures. We find that BipA, a ribosome-associated GTPase, accounts for this temperature-dependent control of biofilm formation by repressing translation of the primary biofilm transcriptional activators VpsR and VpsT at low temperatures. *In vitro* studies demonstrate that low temperatures influence BipA structural conformation and decrease its sensitivity to proteolysis. Proteomic analyses reveal that BipA exerts temperature-dependent control over >200 proteins in *V. cholerae* involved in a multitude of cell processes, including biofilm assembly. Our study reveals a remarkable new facet of the complex *V. cholerae* biofilm regulatory cascade and suggests that combined transcriptional-translational control could be a common mechanism by which bacteria adapt to environmental flux.

## Introduction

A common strategy of bacteria for adaptation and survival to changing environmental conditions is the formation of biofilms: bacterial communities enclosed in an extracellular matrix. *Vibrio cholerae*, the causative agent of the severe diarrhoeal disease cholera, forms biofilms both in biotic and abiotic surfaces in the aquatic environment that inhabits^1-3^ and also in the intestine of the human host^4,5^. The formation of biofilms enhances persistence of *V. cholerae*, since it provides protection against environmental insults, predators and stress conditions and allows a better access to nutrients^3^. There is some evidence indicating that biofilm-formation capacity is critical for intestinal colonization^5^ and biofilms have also been linked to a hyperinfectious phenotype^6^.

*V. cholerae*’s biofilm is primarily composed of VPS (*Vibrio* polysaccharide), matrix proteins (RbmA, RbmC and Bap1) and extracellular DNA^7-13^. The genes encoding the activities for the production of VPS, grouped in the *vpsI* and *vpsII* clusters, and the genes encoding the RbmA and RbmC matrix proteins, located in the *rbm* cluster between *vpsI* and *vpsII* clusters, form the so-called *V. cholerae* biofilm-matrix cluster (VcBMC)^7-10^.

Biofilm formation in *Vibrio cholerae* is a highly regulated process, controlled by the transcriptional activators VpsR, VpsT and AphA, the transcriptional repressors HapR and H-NS, small regulatory RNAs, alternative sigma factors (RpoS, RpoN and RpoE), and small nucleotide signalling (c-di-GMP, cAMP, ppGpp). Specific environmental signals such as changes in salinity, osmolarity, nutrient availability, phosphate limitation, Ca2+ levels, iron availability, and presence of polyamines (spermidine and norspermidine), indole or bile^14,15^ can also affect biofilm formation. Within this complex regulatory network, VpsR and VpsT, whose regulons extensively overlap^16^, are the main transcriptional activators of the *vpsI* and *vpsII* clusters and the *rbmA, rbmC* and *bap1* genes encoding the matrix proteins^17^. HapR is the main repressor of biofilm formation and inhibits transcription of both the activators and the genes encoding the VPS and the matrix proteins^18^. HapR also regulates other processes, such as virulence factor production, type VI secretion^19-21^, and intracellular c-di-GMP levels^18,22^, which in turn indirectly affect biofilm formation. HapR expression is controlled by the quorum sensing cascade that responds to cell density^23,24^, as well as by additional QS-dependent^25-28^ or QS-independent^29,30^ regulators. There is considerable variation in conservation of HapR function between and within the classical (pandemics 1-6), El Tor (pandemic 7) and variant El Tor (late pandemic 7) biotypes that stratify toxigenic *V. cholerae^22,31-33^.*

In both free-living and host-associated environments, *V. cholerae* must adapt to changing extracellular conditions. Specifically, *V. cholerae* experiences a wide range of temperatures, including seasonal and inter-annual temperature changes in the aquatic environment (ranging between 12°C-30°C), and also upon infection of the human host (37°C). Once *V. cholerae* enters the human host, the temperature up-shift controls the expression of virulence factors^34,35^. Adaptation to temperature is important not only for survival in the environment but also for the infection process and for the outbreak and subsequent transmission to a new host.

In this study, we identify VC2744, encoding *V. cholerae* BipA, as a biofilm repressor determinant at low temperatures (i.e. below 22°C). BipA governs a global change in the *V. cholerae* proteome at low temperatures, which includes downregulation of the main biofilm transcriptional regulators VpsR and VpsT. Remarkably, it appears that temperature directly controls BipA activity by altering its structural conformation and turnover. Finally, we show that BipA regulation is independent of HapR, underscoring the intricacies of coordinating transcription and translation processes in response to temperature and cell density to govern biofilm development.

## Results

### Temperature governs colony morphology in *V. cholerae* HapR- strains

Rugose colony morphology in *V. cholerae* is associated with biofilm formation^8^. We noticed that the rugose colony-forming V. *cholerae* El Tor clinical isolate co969 (Figure 1A) lost rugosity below 22°C (Figure 1B), suggesting that temperature had a role in biofilm regulation in this strain. To test whether this phenomenon was due to the higher cell density achieved at higher temperatures, we incubated the plates for longer periods and monitored colony morphology (Figure 1CD). Our results clearly show that the inability of *V. cholerae* co969 to develop rugose colonies at 22°C relies on the temperature, and not population density (Figure 1D).

**Figure 1.**
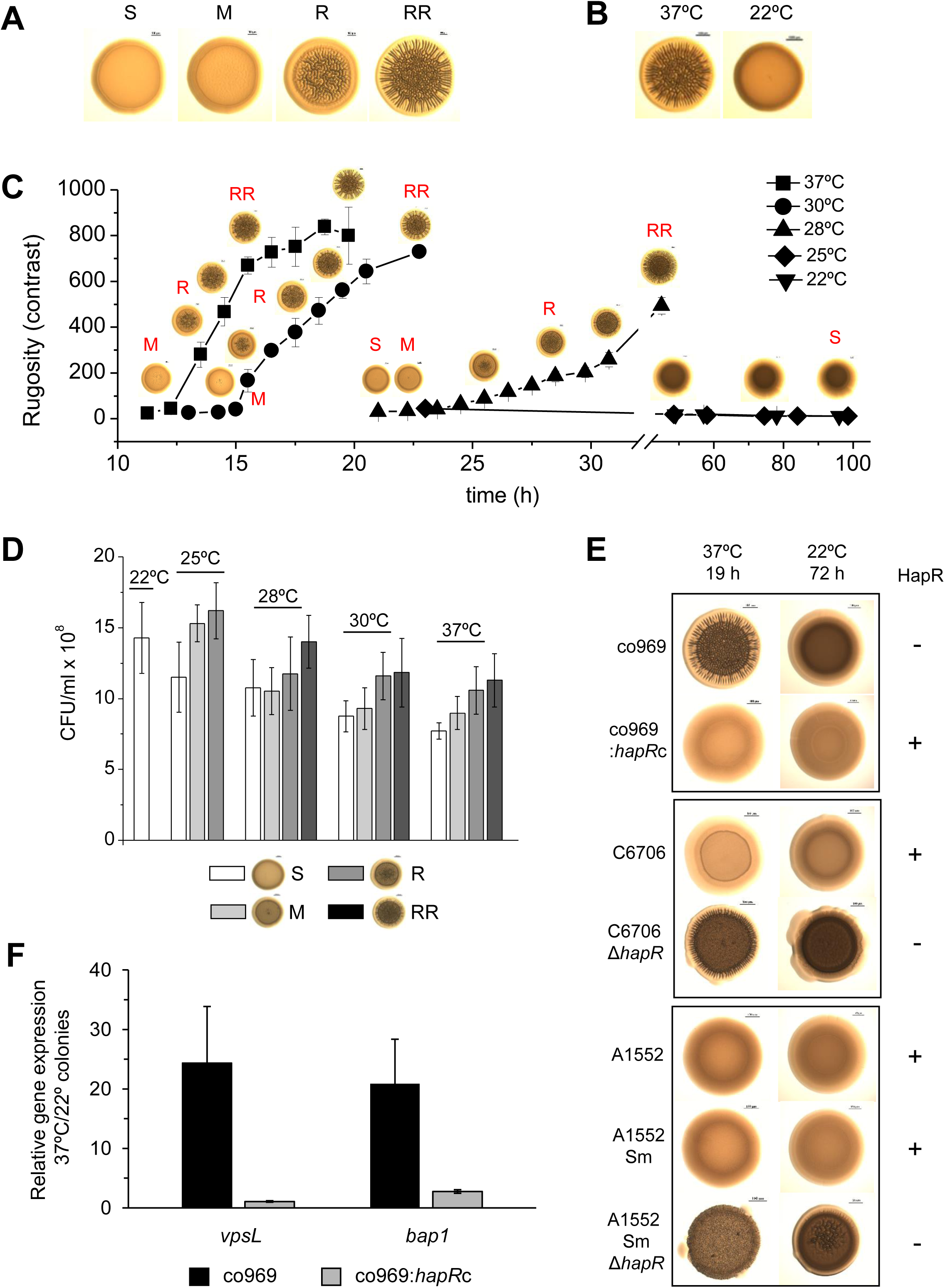
Development of *Vibrio cholerae* co969 colony rugosity is temperature and HapR dependent. **A.** Development of *V. cholerae* co969 colony rugosity at 37°C, divided into four stages: S (smooth colonies), M (transition between smooth and rugose), R (rugose colonies), and RR (very rugose colonies). **B.** *V. cholerae* co969 colony morphology at 37°C and 22°C. **C.** Development of *V. cholerae* co969 colony rugosity over time at different temperatures. Rugosity is represented as contrast calculated using ImageJ software. **D.** Colony forming units (CFUs) per colony for colonies grown at different temperatures and collected at the different rugosity stages shown in the right panel. Values are the average of at least 3 independent experiments having at least 3 biological replicates each. Error bars, standard deviation. **E.** Colony morphology at 37°C and 22°C of co969, co969:*hapR*c (co969 strain carrying the active variant of *hapR* from C6706, *hapR*c), and C6706 and A1552 and their respective Δ*hapR* mutant derivatives. The used incubation times at different temperatures were previously optimized to result in colonies with a comparable number of CFU per colony. **F.** Relative expression of *vpsL* and *bap1* between colonies grown at 37°C vs 22°C for co969 and co969:*hapR*c strains, determined by qRT-PCR. qRT-PCR results are the average of 3 biological replicates, each replicate containing 8 colonies; error bars, standard deviation. Expression of *hfq* was used as a control.

*V. cholerae* co969 naturally lacks wild-type (WT) HapR due to a frame shift mutation that results in a truncated version of the protein. We observed that, similarly to co969, other *V. cholerae* strains with inactive HapR variants also developed rugosity in a similar temperature-dependent fashion both in solid-air (i.e. colony morphology) and air-liquid interphases (i.e. wrinkled pellicles) (SM Figure 1). However, we did not observe this phenotype in HapR+ *V. cholerae* strains such as C6706 and A1552 (SM Figure 1), which were smooth at all temperatures. Remarkably, deletion of *hapR* in these strains led to temperature-dependent rugosity development comparable to the naturally *hapR*-deficient strains, and substitution of the mutated *hapR* locus in co969 with the WT *hapR* copy from C6706 (*hapR*c) rendered smooth colonies at all temperatures (Figure 1E). Therefore, the temperature-dependent biofilm control in *V. cholerae* is HapR-independent, and is masked in HapR+ strains.

We next measured the expression of specific structural biofilm components at different temperatures using qRT-PCR (Fig 1F). Both *vpsL* and *bap1* were up-regulated in co969 colonies grown at 37°C vs 22°C (as in Figure 1B), indicating that colony rugosity can be used as a readout to study temperature-dependent biofilm development regulation in *V. cholerae*. Furthermore, *vpsL* and *bap1* expression did not differ between these two temperatures in the co969:*hapR*c background, buttressing the idea that the temperature-dependent program governing biofilm development in *V. cholerae* is masked by HapR (Figure 1F).

### Biofilm genes are upregulated in colonies grown at 37°C vs 22°C

As *V. cholerae* biofilm formation is largely transcriptionally controlled, to understand how temperature influenced the global *V. cholerae* biofilm regulon, we performed RNA-seq of *V. cholerae* co969 WT colonies grown at 37°C (rugose, 37R) vs 22°C (smooth, 22Sb) (Figure 2). As controls, we performed transcriptomic analyses of smooth colonies from an earlier point (time 1, SM Figure 2) grown at 37°C (37S) and 22°C (22S) to eliminate temperature-independent or cell-density driven gene expression changes. In agreement with our phenotypic observations, expression of the *vpsI* and *vpsII* gene clusters (e.g. *vpsL*), encoding the activities responsible for the production of the exopolysaccharide (VPS), and the *rbmA, rbmC* and *bap1* genes, encoding the biofilm matrix proteins, were highly upregulated (ca. 10-20 fold) in 37R compared to 22Sb colonies (Figure 2C, SM Figure 2 and SM Tables 4, 5 and 6). These results were validated by qRT-PCR (Figure 2D). Interestingly, most known biofilm transcriptional regulators (e.g. *vpsT, vpsR, hapR*), as well as other genes involved in quorum sensing, type II secretion, and c-di-GMP signalling were not differentially expressed (SM Figure 2; SM Tables 4, 5 and 6). These results suggested that temperature-dependent biofilm formation is controlled by either an unknown transcriptional regulator or a post-transcriptional mechanism.

**Figure 2.**
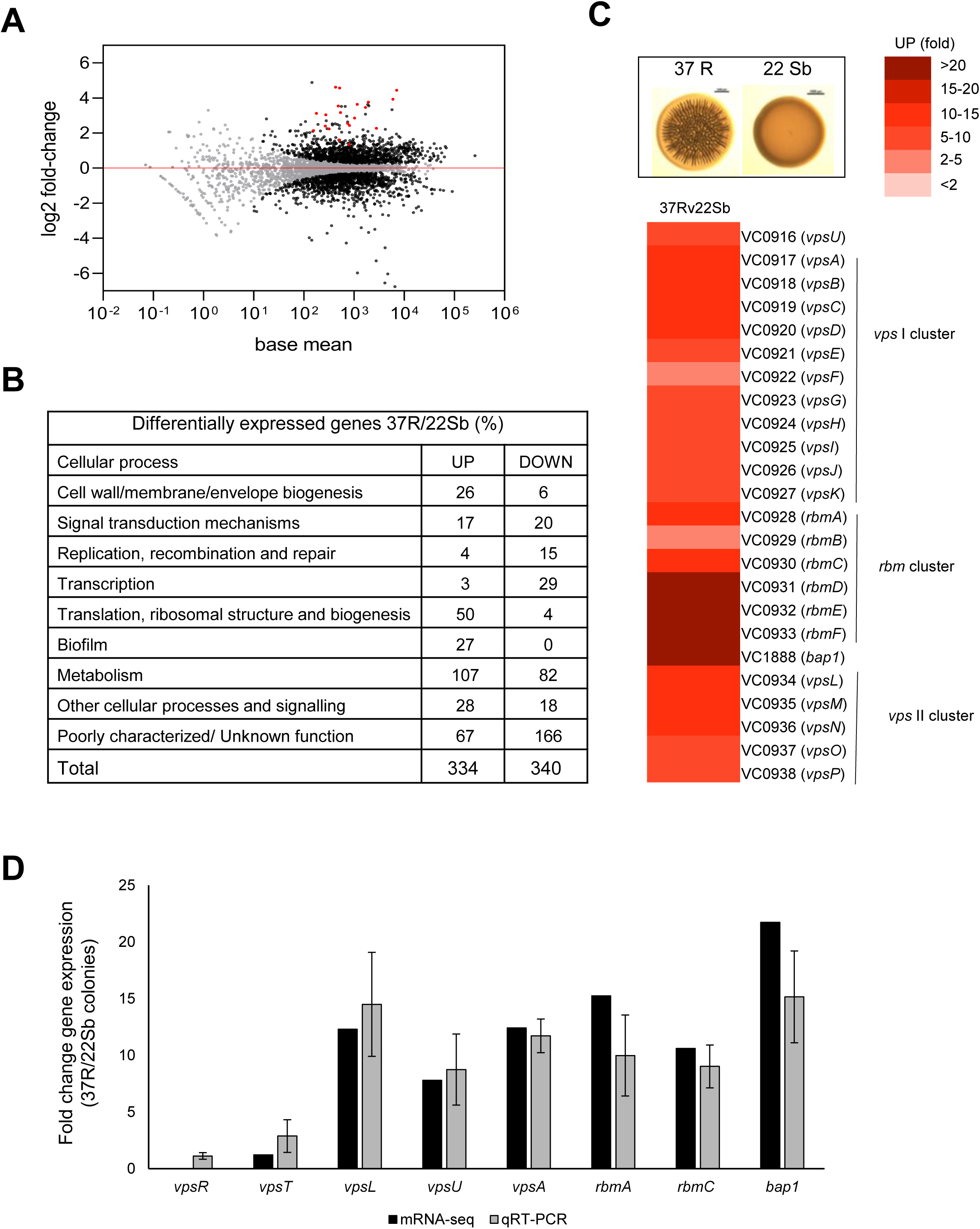
Biofilm components are transcriptionally upregulated at higher temperatures. Comparison of relative gene expression between *V. cholerae* co969 colonies incubated at 37°C (rugose, 37R) and 22°C (smooth, 22Sb) by mRNA-seq. Rugose colonies collected at 37°C and smooth colonies collected at 22°C had a similar number of CFUs per colony, as described under experimental procedures. **A.** MA plots representing the log2FC against mean expression for differentially expressed genes between 37R vs 22Sb colonies. Black dots, significantly differentially expressed genes; grey dots, not significantly differentially expressed genes; red dots, genes encoding biofilm structural components (*vpsI* and *vpsII* clusters, *rbmA, rbmC* and *bap1)*. **B.** Table showing the number of differentially expressed genes (UP and DOWN-regulated) between 37R vs 22Sb colonies, grouped by COG categories. **C.** Heat map showing the differential relative expression (fold change) between 37R and 22Sb of genes belonging to the *vpsI, vpsII* and *rbm* clusters and *bap1*, encoding the VPS (*Vibrio* exopolysaccharide) and biofilm matrix proteins (RbmA, RbmC and Bap1), respectively. **D.** Validation of differential gene expression obtained by mRNA-seq by qRT-PCR. The graph represents a comparison of the relative gene expression - obtained in both the mRNA-seq analysis and qRT-PCR experiments - of biofilm activators (*vpsR* and *vpsT*), genes encoding VPS (*vpsL, vpsA* and *vpsU*) and matrix proteins (*rbmA*, rbmC and *bap1*) between 37R and 22Sb colonies. Values are the average of 3 independent qRT-PCR experiments containing 3 biological replicates each, each replicate containing 8 colonies pooled together, and 3 technical triplicates of each biological replicate; error bars, standard deviation. Expression of *gyrA* was used as a control.

### A genetic screen for determinants of temperature-dependent colony rugosity

We reasoned that our observations could stem from either activation (at 37°C) and/or repression (at 22°C) of gene expression. Since more is known about biofilm regulation at 37°C^14^ and mutagenesis at this temperature would produce a high number of false positives (i.e. all known structural biofilm proteins and temperature-independent activators), we decided to perform a loss-of-function screen for potential biofilm repressors at low temperatures. We performed random transposon mutagenesis in *V. cholerae* co969 and selected mutants that inappropriately formed rugose colonies at 22°C (Figure 3A). Out of 11,000 mutants screened, 459 colonies had a rugose phenotype at 22°C. Pooled sequencing of the 459 rugose colonies identified insertions in 99 genes, corresponding to diverse cell functions such as flagellar motility, chemotaxis, c-di-GMP signalling, LPS, transport, cell wall, PTS systems, regulatory functions, transport, translation and others (Figure 3B, SM Table 6). To filter out potential temperature-independent repressors, we performed a similar screen for rugose colonies at 37°C using the temperature-insensitive, smooth *V. cholerae* strain C6706 (SM Table 7) and removed hits from this screen from our list. After filtering, the most frequently inserted gene in co969 rugose colonies at 22°C was VC2744 (see SM Table 6), a protein with a 74% identity to *Salmonella enterica* serovar Typhimurium BipA, also referred to as TypA in other organisms^36,37^. BipA has been implicated in the regulation of virulence and stress responses in bacteria^38-43^. Mutations in BipA have been associated with a cold-sensitive phenotype in *E. coli*^44^, but a clean deletion of *vc2744* in *V. cholerae* co969 did not influence growth or cell morphology at any temperature (SM Figure 3). In other bacteria, BipA has been implicated in biofilm formation^41,45,38,46^, but the mechanism behind this phenotype has remained elusive. As BipA is thought to be a ribosome-binding GTPase that regulates ribosome assembly^47^, we reasoned that VC2744 could be repressing biofilm development at low temperatures through translational regulation.

**Figure 3.**
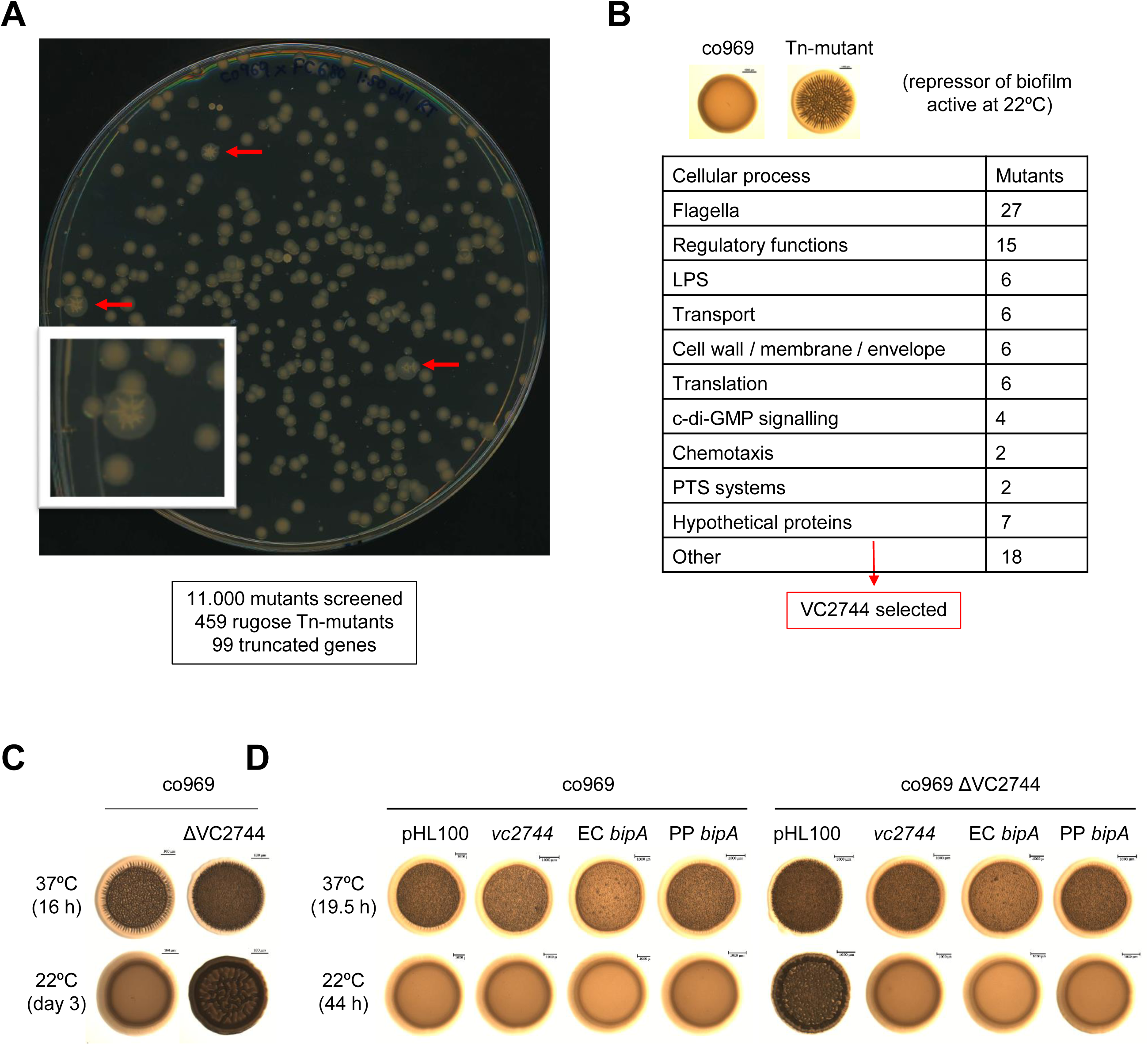
Transposon mutagenesis of *V. cholerae* at 22°C identified VC2744 as a biofilm repressor. **A.** Example of an LB (10 g/l NaCl) + Sm200 + Km50 agar plate used for selection of *V. cholerae* co969 transposon mutants with a rugose colony phenotype (pointed with red arrows) at 22°C. The number of screened transposon mutants, rugose colonies selected and final number of truncated genes is indicated below. **B.** Table indicating the number of truncated genes identified grouped into different categories. Out of these genes, VC2744 - encoding *bipA* - was selected due to its high representation (see SM Table 6). **C and D.** Effect of VC2744 on co969 colony morphology. **C.** Deletion of VC2744 on co969 results in a rugose colony phenotype at 22°C. **D.** Colony morphology at 37°C and 22°C of co969 and co969 ΔVC2744 carrying either pHL100- *bipA*, for overexpression of *V. cholerae* co969 *vc2744* from the IPTG-inducible P*lac* promoter, pHL100-EC*bipA* or pHL100-PP*bipA*, for overexpression of the *bipA* variants of *E. coli* MG1655 K-12 or *P. putida* KT2440, respectively, or the empty plasmid (pHL100). Overexpression of the different BipA variants restored the smooth colony phenotype at 22°C, while it only resulted in slightly decreased rugosity at 37°C.

### BipA represses biofilm development at low temperature

Deletion of *vc2744* in co969 confirmed the rugose colony phenotype from our screen at 22°C (Figure 3C). Complementation of co969 Δ*vc2744* with either *V. cholerae vc2744* or *bipA* orthologues from *E. coli* MG1655 K-12 or *Pseudomonas putida* KT2440 led to restoration of the WT smooth colony morphology phenotype at 22°C (Figure 3D). These results, together with the high protein similarity between orthologues, strongly suggest that *vc2744* encodes a *bipA* homolog in *V. cholerae*. Consistent with our previous data, deletion of *bipA* led to an increase in rugosity only in HapR- (co969 or C6706 Δ*hapR*) but not HapR+ strains (C6706 wt or co969:*hapR*c), suggesting that the effect of BipA on colony morphology is epistatic to HapR in certain *V. cholerae* strains (SM Figure 4A and B).

**Figure 4.**
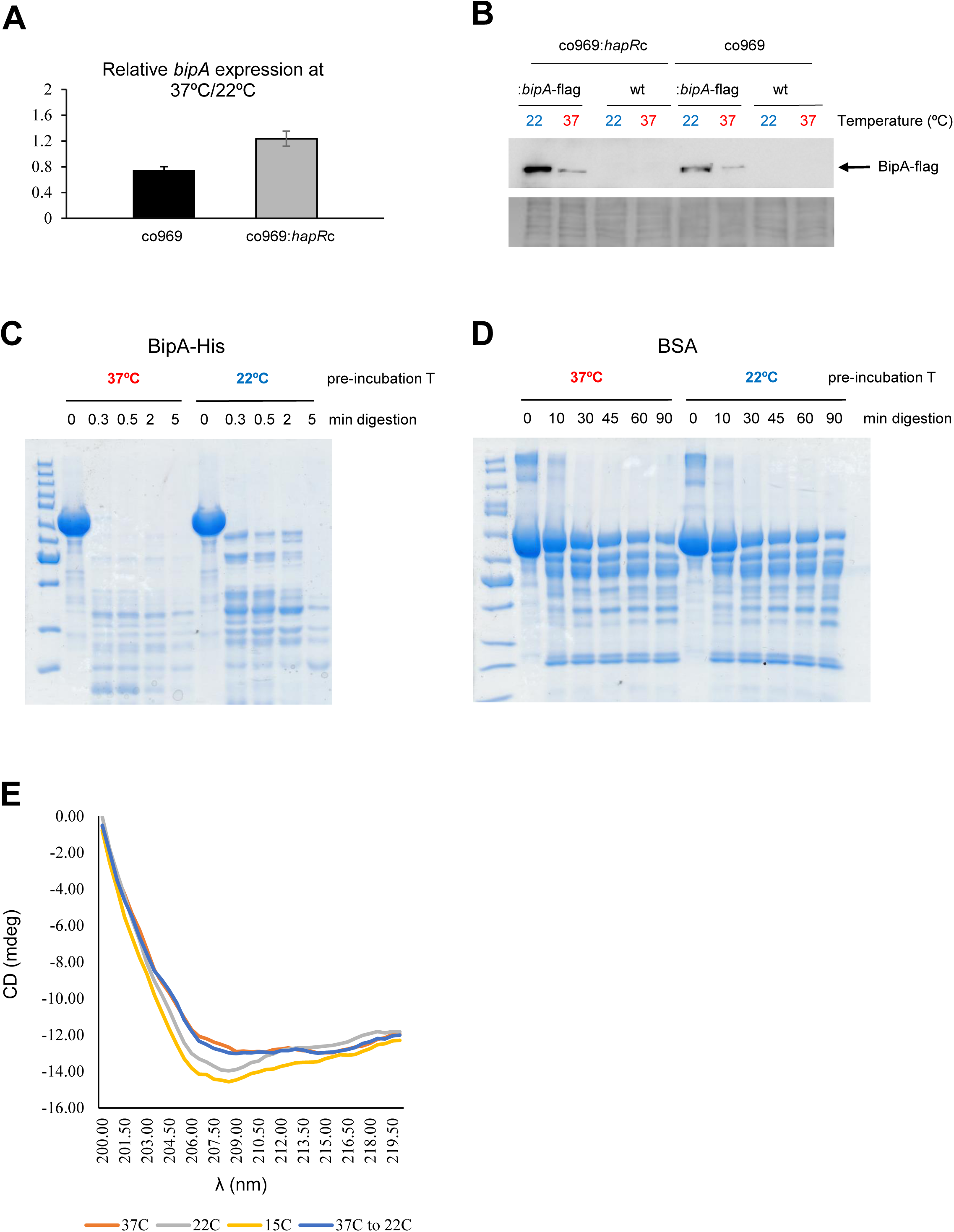
BipA protein levels are elevated at 22°C. **A.** Relative *bipA* expression in *V. cholerae* co969:*bipA*-flag or co969:*hapR*c:*bipA*-flag strains, carrying a chromosmal *bipA*-flag phusion, between rugose colonies grown at 37°C (37R) and smooth colonies grown at 22°C (22Sb), determined by qRT-PCR. Data are the average of 3 biological replicates, each containing 8 colonies pooled together. Error bars, standard deviation. **B.** Western blot showing the BipA-flag protein levels of co969:*bipA*-flag or co969:*hapR*c:*bipA*-flag rugose colonies grown at 37°C (37R) and smooth colonies grown at 22°C (22Sb). co969 and co969:*hapR*c strains were used as a negative controls, respectively. The gel is a representative of 5 independent experiments. The Coomassie staining of the membrane, used as control, is shown below. **C.** Sensitivity to partial digestion with trypsin of purified BipA-His pre-incubated at either 37°C or 22°C. **D.** Sensitivity to partial digestion with trypsin of BSA pre-incubated at either 37°C or 22°C. In both **C** and **D**, 25 μg aliquots of BipA-His or BSA, respectively, were pre-incubated at either 37°C or 22°C for 2 h; then, they were kept at room temperature for 5 min and subjected to digestion with 1.5 μg trypsin at room temperature during the incubated times: 0, 20 sec, 30 sec, 2 min and 5 min, for BipA-His; or 0, 10, 30, 45, 60 and 90 min, for BSA. The digested fragments were separated on a 10% acrylamide gel, and the gel was stained with Coomassie. **E.** Circular dichroism curve (mdeg represented against wavelength) of purified BipA-His pre-incubated at different temperatures (37°C, 22°C or 15°C) for 30 min, or pre-incubated at 37°C for 30 min and then sifted to 22°C and further incubated for 30 min.

A previous *V. cholerae* transcriptomic survey indicated that a mutant in the biofilm regulator *vqmA* increased *bipA* levels^29^. However, our results clearly show that deletion of *vqmA* did not phenocopy Δ*bipA* rugose colony morphology at low temperature, suggesting that VqmA does not affect expression of *bipA* under these biofilm-inducing experimental conditions (SM Figure 4C). To learn more about the regulatory network of BipA, we deleted this gene in the Δ*vpsR* and Δ*vpsT* backgrounds, which have constitutively smooth colony morphologies. Both double mutants maintained smoothness (SM Figure 4C), suggesting that BipA acts upstream of VpsR and VpsT. Collectively, our results demonstrate that BipA is a temperature-dependent, HapR-independent repressor of *V. cholerae* biofilm formation.

### BipA stability is controlled by temperature

A key observation was that *bipA* overexpression at 37°C rendered only a very subtle reduction in rugosity compared to that at 22°C (Figure 3C), suggesting that BipA activity is higher at lower temperatures. Remarkably, even though *bipA* expression was largely unchanged between colonies grown at 37°C and 22°C (SM Table 4 and Figure 4A), BipA protein levels were about 10 times higher at 22°C (Figure 4B). Temperature-dependent changes in BipA levels were independent of HapR status (Figure 4A and B), underscoring the idea that these two proteins use distinct mechanisms to repress biofilm formation.

We next performed circular dichroism studies to study whether BipA could be directly structurally regulated by temperature. Analysis of purified BipA pre-incubated at different temperatures (37°C, 22°C, and 15°C) suggested differences in protein folding between these temperatures (Figure 4E). Interestingly, incubation of BipA at 37°C followed by subsequent incubation at 22°C showed the same pattern as for the protein incubated at 37°C (Figure 4E) suggesting that these changes are irreversible. To determine if these differences in conformation rendered BipA more susceptible to degradation at higher temperatures, we incubated purified BipA either at 37°C or 22°C and then checked their susceptibility to trypsin-mediated limited proteolysis. BipA incubated at 37°C was degraded markedly faster than protein incubated at 22°C (Figure 4C). We did not observe such temperature-dependent proteolytic sensitivity for the control protein BSA (Figure 4D), suggesting that specific and direct temperature control of BipA folding promotes increased turnover at higher temperatures.

As these data suggested a scenario where BipA is more susceptible to degradation at 37°C, we aimed to identify the BipA-targeting protease. A transposon mutagenesis screening for aberrantly smooth co969 colonies at 37°C identified the LonA protease as a potential repressor of BipA activity (SM Figure 5B and SM Table 8). LonA has been previously reported to have a role in biofilm regulation^48^. Deletion of VC1920 (*lonA*) in co969 confirmed formation of smooth colonies at both 37°C and 22°C (SM Figure 5B) and overexpression of *lonA* resulted in increased rugosity at both temperatures (SM Figure 5C). However, BipA protein levels were unchanged in the Δ*lonA* background (SM Figure 5D), suggesting that the effect of LonA in biofilm formation does not involve BipA. Accordingly deletion of *lonA* in the co969 Δ*bipA* background also resulted in smooth colonies at all temperatures, and overexpression of *lonA* in co969 Δ*bipA* further increased rugosity (SM Figure 5B and C). Further research will be needed to clarify LonA’s role in biofilm formation and to identify the putative protease controlling BipA turnover in *V. cholerae*.

**Figure 5.**
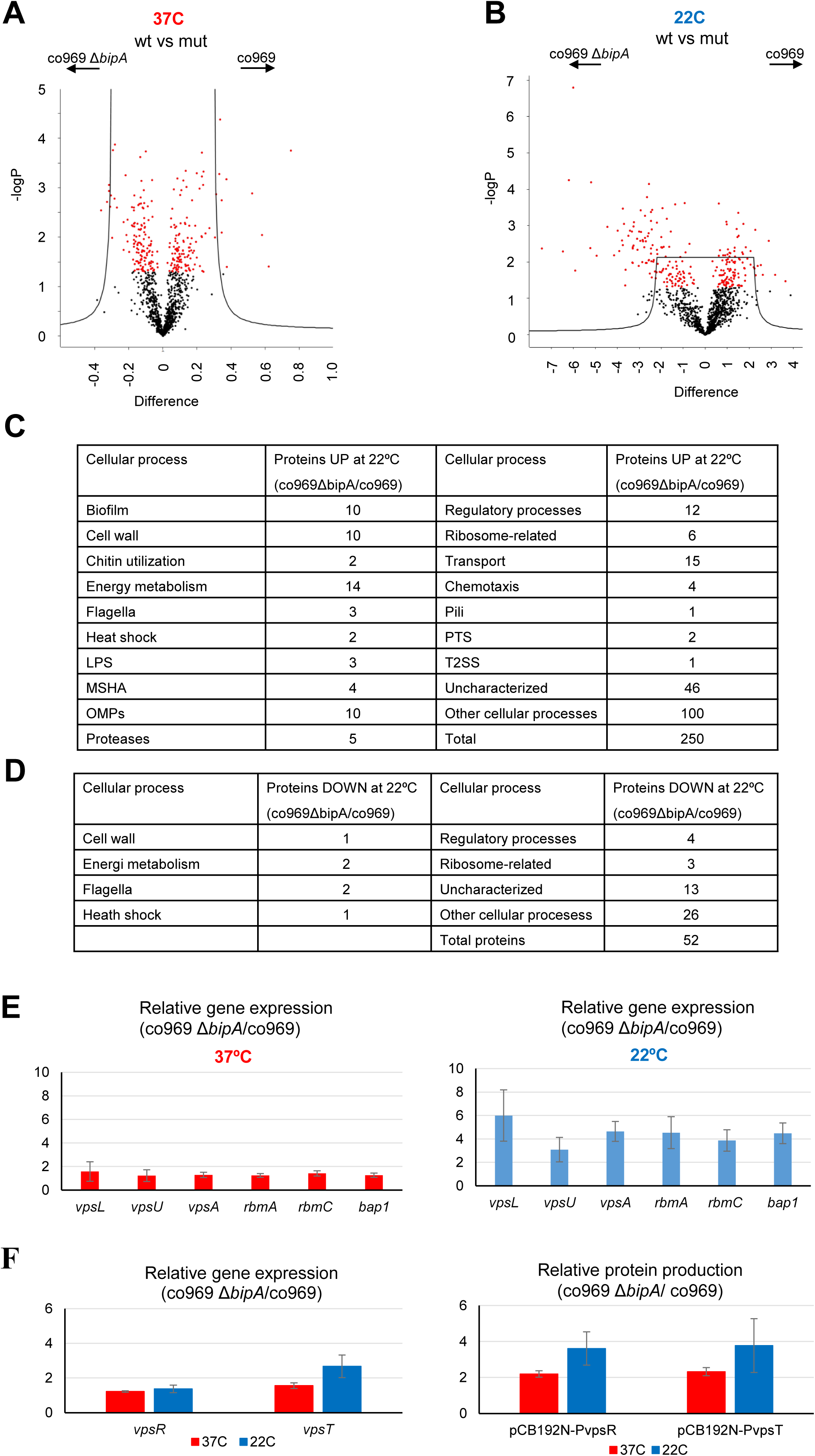
Global proteomic analysis of *V. cholerae* co969 wt vs Δ*bipA* colonies grown at 37°C and 22°C. **A and B.** Volcano plots representing the log t-test P-value against the t-test difference for a comparison of protein levels between co969 wt vs Δ*bipA* colonies grown at either 37°C (**A**) or 22°C (**B**). Black dots represent proteins belonging to the pool of non-differentially regulated proteins between the wt and the Δ*bipA* strain; red dots represent proteins that are significantly regulated between both strains at the given condition. Higher numbers of t-test difference and log t-test P-value indicate more differentially regulated proteins. **C and D.** Table indicating the number of differentially expressed proteins between co969 wt vs Δ*bipA* colonies at 37°C (**C**) and 22°C (**D**) grouped by function. **E.** Relative expression between co969 Δ*bipA* and co969 strains of genes encoding the exopolysaccharide (*vpsA, vpsL, vpsU*) and the biofilm matrix proteins (*rbmA, rbmC, bap1)* in rugose colonies grown at 37°C (left panel) or smooth colonies grown at 22°C (right panel), determined by qRT-PCR. **F** Left graph, relative expression of *vpsR* and *vpsT* genes encoding the biofilm activators between co969 Δ*bipA* and co969 strains in rugose colonies grown at 37°C (left) or smooth colonies grown at 22°C (right), determined by qRT-PCR. Right graph, relative Miller activity between co969 Δ*bipA* and co969 strains carrying the translational β-galactosidase reporters pCB192N-P_*vpsR*_ or pCB192N-P_*vpsT*_, in rugose colonies grown at 37°C or smooth colonies grown at 22°C, determined by β-galactosidase assays. In both **E** and **F**, qRT-PCR values are the average of 2-3 independent experiments containing 3 biological replicates each (each replicate containing 8 colonies); error bars, standard deviation. Expression of *hfq* was used as a control. In **F**, β-galactosidase assays results are the average of 3-4 independent experiments containing 3 biological replicates each (each replicate containing 4 colonies); error bars, standard deviation.

### BipA controls translation at lower temperatures

As BipA is a ribosome assembly factor at low temperatures^47,49^, we wondered if it exerted a global effect on translation along with its role in controlling biofilm formation. To address this question, we performed global proteomic analyses of co969 WT and Δ*bipA* grown at either 37°C (rugose colonies) or at 22°C (smooth colonies), followed by 4 comparative analyses: (i) WT vs Δ*bipA* at 37°C; (ii) WT vs Δ*bipA* at 22°C colonies; (iii) WT 37°C vs 22°C colonies; and (iv) Δ*bipA* 37°C vs 22°C colonies. The proteomic analysis was able to identify 1639 (43%) of 3783 known *V. cholerae* proteins, out of which 695 were significantly differentially abundant.

The difference between co969 WT and Δ*bipA* proteomes was greater at 22°C than at 37°C, further suggesting that BipA activity is highest at lower temperatures (Figure 5 A and B). Relative to the WT strain, 250 proteins were up-regulated in Δ*bipA* at 22°C, and 52 proteins were down-regulated (Figure 5B). The most represented differentially regulated proteins between co969 WT and Δ*bipA* at 22°C were related to external cellular components and biological processes involved in localization and transport (Figure 5B and C). We identified a number of biofilm components among the up-regulated proteins, confirming the role of BipA as a biofilm repressor at low temperatures (i.e. 22°C). Among the down-regulated proteins, we found flagella components. Consistent with this finding, the co969 Δ*bipA* mutant exhibited 10-20% reduced motility than the WT strain, with motility slightly more reduced at 22°C than at 37°C (SM Figure 6A). Interestingly, a similar scenario occurred in the co969:*hapR*c background (SM Figure 6C), indicating that the effect of BipA on motility is independent of the presence of HapR. Complementation of co969 Δ*bipA* at 22°C, but not at 37°C, almost completely restored the WT motility phenotype (SM Figure 6B) further strengthening the role of BipA at lower temperatures.

**Figure 6.**
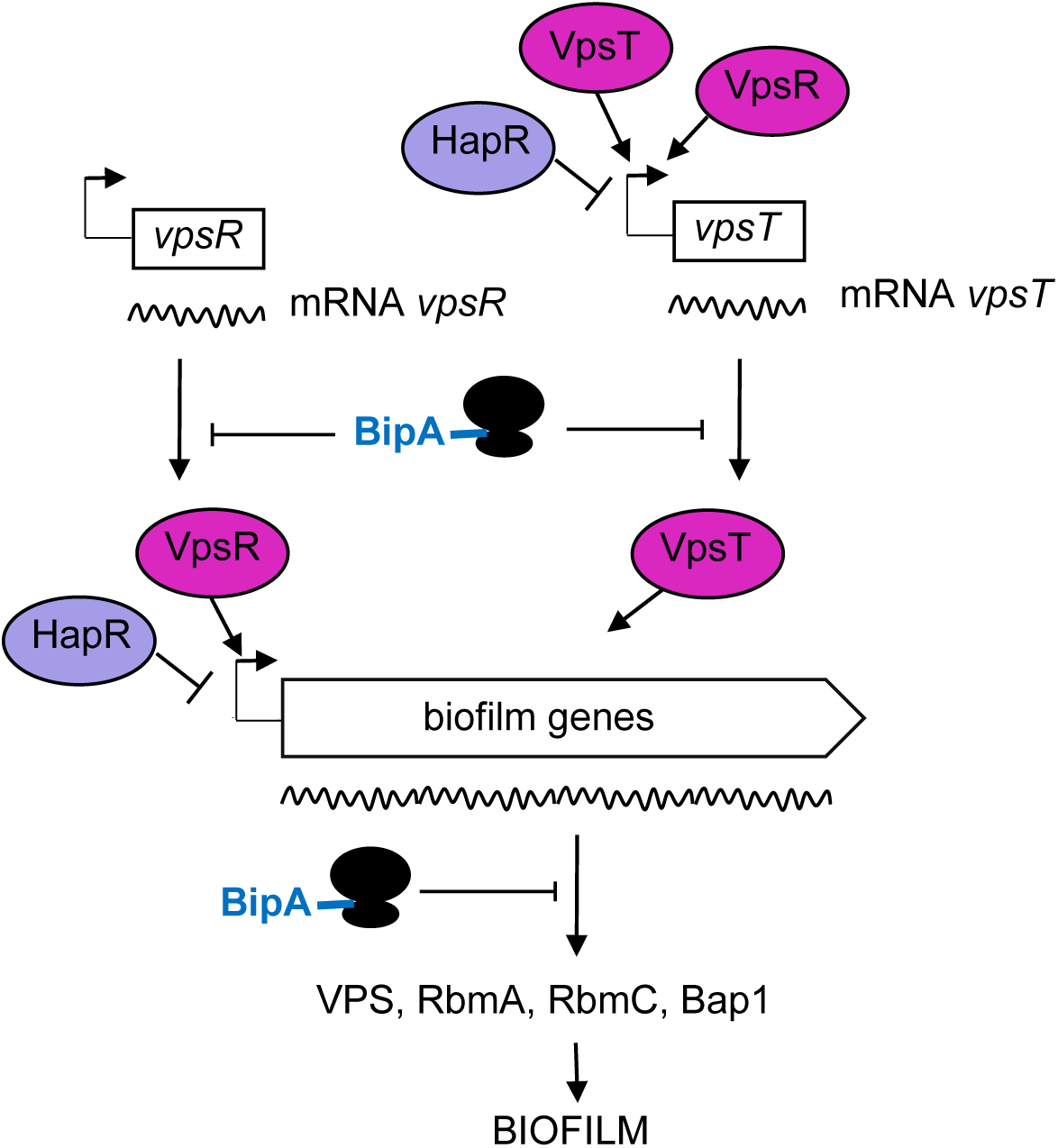
Proposed model for BipA within the *V. cholerae* biofilm regulatory cascade. Schematic showing the proposed model for the interplay between the main biofilm transcriptional regulators VpsR, VpsT and HapR, and the translational repressor BipA. In HapR- strains transcription of *vpsR* and *vpsT* and the biofilm genes (*vpsI* and *vpsII* clusters encoding the VPS and *rbmA, rmbC* and *bap1* encoding the matrix proteins) leads to biofilm formation at 37°C but not at 22°C, when high levels of BipA inhibit translation of the mRNAs of both the activators and the biofilm genes. In HapR+ strains transcription of both the biofilm activators and the biofilm genes is negatively regulated by HapR, and no biofilm is be produced. At 22°C BipA confers an extra check point in this regulation by making sure that, even if residual levels of mRNA would be produced, their translation would then be inhibited, resulting in no biofilm formation.

### BipA inhibits VpsT and VpsR translation at low temperatures

The proteomic analysis revealed that VpsR and VpsT protein levels were higher at 22°C in the co969 Δ*bipA* background relative to the WT and higher at 37°C than at 22°C for the WT background (SM Table 9). In contrast, our RNAseq data and targeted qRT-PCR studies indicated that the expression of the two major biofilm transcriptional regulators *vpsR* and *vpsT* was largely unchanged at different temperatures and between Δ*bipA* and WT cultures (Figure 5F and SM Tables 4, 5 and 6). Transcription of known VpsR and VpsT targets (*vpsU, vpsA, vpsL, rbmA, rbmC* and *bap1*) in co969 Δ*bipA* vs co969 colonies at different temperatures tracked with changes in VpsR and VpsT protein levels (Figure 5E). These observations suggested that BipA could negatively regulate translation of VpsR and VpsT at lower temperatures, indirectly repressing expression of the *V. cholerae* biofilm regulon.

Finally, to directly assess the effect of BipA in controlling VpsT and VpsR translation, we performed β-galactosidase assays with co969 Δ*bipA* and co969 strains carrying the translational reporter vectors pCB192N-P_*vpsT*_ and pCB192N-P_*vpsR*_. The results showed that translation of both activators was upregulated in the co969 Δ*bipA* mutant vs co969 WT, especially at 22°C (Figure 5F, right graph). Collectively, these results support a role for BipA as a negative regulator of biofilm formation through the specific control of VpsR and VpsT protein levels.

## Discussion

Although *V. cholerae* biofilms are a key component of this pathogen’s success, the molecular mechanisms that underlie this process are complex and rely on many different inputs and pathways that converge on the regulation of the core biofilm regulon. Previous studies described the existence of a complex interplay of regulatory mechanisms that ultimately result in the transcriptional control of the biofilm genes. Here, complementary genetic and proteomic approaches led us to discover that BipA controls temperature-dependent changes in biofilm formation in *V. cholerae* HapR- strains via post-transcriptional regulation of the master biofilm regulators VpsT and VpsR. Remarkably, BipA protein stability is enhanced at low temperatures, allowing this protein to control the levels of >200 proteins in *V. cholerae*, and indicating that outside of its effect on biofilm formation, BipA may shape other aspects of pathogen physiology at low temperatures.

BipA is thought to act as a ribosome assembly factor^47,50,49^ that modulates translation of a specific and distinct subset of mRNAs under specific stress conditions^50^. To explain this, it has been proposed that BipA together with ppGpp controls ribosome binding specificity to regulate the synthesis of virulence or stress factors^40^. However, other studies suggest that BipA is important for biogenesis of the 50S^49,51,52^. Contrary to the cold sensitive phenotype of *E. coli bipA* mutant^44^, growth of *V. cholerae* co969 Δ*bipA* was not affected at 22°C (SM Figure 3), suggesting that impact of BipA on biofilm formation is more likely due to a re-organization of ribosomal translation specificity rather than a general defect in ribosome assembly and protein synthesis.

Our results demonstrate that BipA represses biofilm at lower temperatures by regulating the synthesis of the main biofilm regulators and structural components. Outside of a direct effect on the translation of VpsT and VpsR, it is possible that BipA also indirectly represses biofilm development by promoting the synthesis of flagellar components (SM Figure 6), since it is known that bacterial motility and biofilm formation are inversely correlated^30,53^. Finally, we cannot exclude that BipA further regulates translation of other activities involved in biofilm formation, such as diguanylate cyclases (DGCs) or phosphodiesterases (PDEs) controlling c-di-GMP levels. Further research will help to untangle the biofilm regulatory network controlled by BipA.

Our observations of biofilm repression at low temperatures conflict with results previously reported for *V. cholerae* A1552 and *V. salmonicida*, which both encode functional HapR homologs. In *V. cholerae* A1552, increased c-di-GMP production at low temperatures by specific DGCs and the cold-shock gene *cspV* leads to higher biofilm formation^54^ ^55^. Since it is known that *V. cholerae* El Tor and classical biotypes (HapR+ and HapR-, respectively) modulate c-di-GMP levels using different pathways^22^, which in turn results in differential regulation of biofilm formation, it seems likely that the presence or absence of HapR could account for the observed differences in the temperature-dependent regulation between these *V. cholerae* strains. Indeed, the 6 DGCs that were induced at low temperature in *V. cholerae* A1552, and account for the increased biofilm production at low temperature, did not appear to be differentially expressed in our RNA-seq studies of co969. These results further underscore the diversity of biofilm regulatory processes in pathogenic *V. cholerae*, especially in strains where the master regulator HapR is absent.

Even though the temperature-regulatory effect on *V. cholerae* colony morphology was seemingly masked by HapR (SM Figure 1 and Figure 1E), we showed that HapR and BipA function independently from each other (SM Figure 4 A and B). HapR does not control BipA expression (Figure 4A and B) and it does not affect the approximate 20% reduction of motility between co969 wt and Δ*bipA* (SM Figure 6, compare panels A and C). Although HapR or BipA alone (HapR+ strains and high temperature or HapR-strains at low temperature, respectively) are sufficient to prevent rugose colony morphology (SM Figure 4, panels A and B), these proteins act together to ensure biofilm repression through both transcriptional and translational control. Moreover, our proteomics data showed that BipA, like HapR, is a global regulator that may impact manifold processes such as virulence, competence, quorum sensing and the T6SS. We propose that biofilms represent just one example of a cellular pathway whose regulation involves combined transcriptional-translational control mechanisms.

Since BipA is highly conserved amongst bacteria^36,37,49^ it is tempting to speculate that the BipA-dependent control of biofilm formation by temperature observed in *V. cholerae* could be conserved in other bacteria. In agreement with this hypothesis, deletion of BipA has been associated to increased biofilm levels in *P. aeruginosa* PAO1^41^ and *Bordetella holmesii*^45^. BipA from *E. coli* and *P. aeruginosa* cross-complemented *V. cholerae ΔbipA* temperature-dependent phenotypes (Figure 3D), further supporting a model of BipA function as a bacterial “temperature sensor”.

Since bacteria thrive within a wide range of temperatures, it would be interesting to identify the amino acid residues in the sequence of BipA that dictate its capacity to change conformation, stability and half-life in response to temperature fluctuation. Differences in the sequence of BipA may also explain distinct regulatory outcomes between species. For example, BipA is crucial for resistance to the antimicrobial peptide P2 in *E. coli* and *Salmonella* spp.^56,57^, but not in *V. cholerae*^58^, whose BipA sequences are not identical.

Collectively, defining the cellular processes and regulatory networks that lay under the control of BipA will shed light on how *V. cholerae* coordinates multiple behaviours in response to temperature changes. Future research will address the effect of temperature-dependent BipA regulation in biofilm dispersal during host-environment transitions and the associated potential consequences in cholera transmission and outbreaks.

## Materials and Methods

### Colony morphology assays

2 μl drops of *V. cholerae* o/d cultures, started from a 1:100 dilution of an o/n culture, were dispensed in LB agar plates containing 10 g/l NaCl and incubated the appropriate time at the mentioned temperature (37, 30, 28, 25 or 22°C). The morphology of the colonies was recorded using a Nikon SMZ1500 stereomicroscope.

### Pellicle biofilm assays

5 μl drops of *V. cholerae* o/d cultures, started from a 1:100 dilution of an o/n culture, were dispensed into 12 well plates, each well containing 5 ml of LB (10 g/l NaCl), and incubated the appropriate time at the mentioned temperature (37, 30, 28, 25 or 22°C). Images of pellicle biofilms were recorded using a Nikon SMZ1500 stereomicroscope.

### Motility assays

2 μl drops of *V. cholerae* o/n cultures were dispensed in 0.3% LB agar plates containing 10g/l NaCl and incubated the appropriate time at 37 or 22°C. For *V. cholerae* strains carrying pHL100-derivatives, 0.3% LB agar plates containing 10g/l NaCl + Km (50 μg/ml) + IPTG (isopropyl-β-d-thiogalactosidase, 1 mM) were used. Motility was determined by measuring the ratio using ImageJ^59^.

### RNA isolation

2 μl drops of *V. cholerae* o/d cultures, started from a 1:100 dilution of an o/n culture, were dispensed in LB agar plates containing 10 g/l NaCl and incubated the appropriate time at either 37or 22°C. For each sample, 8 colonies were collected, pooled together in an Eppendorf tube for RNA extraction and stored at −20°C. For comparative analysis, the OD_600_ of the resuspended colonies, as well as their CFU/colony and total protein content were similar for the colonies grown at 37°C and 22°C at the selected sample collection time points (time 1 and time 2), even though the incubation time was different. Pellets were then resuspended in 400 μl of solution containing 10% glucose, 12.5 mM Tris pH 7.6 and 5 mM EDTA and, after adding 60 μl 0.5 M EDTA, cells were disrupted on a bead beater (4°C, maximum speed for 1 min 15 sec) in the presence of 0.5 ml of acid phenol. After centrifugation, 1 ml Trizol (Ambion) was added to the supernatant, incubated at room temperature for 10 min, 100 μl choloroform:IAA were added, mixed by vortex for 10 sec, and centrifuged at 14.000 rpm 4°C for 15 min. After two more chloroform:IAA extractions, the RNA was precipitated by mixing the aqueous phase with 0.7 volumes isopropanol, incubated for 30 min at −20°C, centrifuged at 14.000 rpm 4°C for 30 min, and washed with 70 % ethanol. RNA pellets were then resuspended in 200 μl H_2_O and subjected to two consecutive DNaseI (Roche)-treatments for 1h at 37°C, in the presence of RNase inhibitor. After a phenol:chloroform:IAA and a chloroform:IAA extractions, the RNA was precipitated by mixing the aqueous phase with 3 volumes of 95% ethanol and 1/10 volumes of 3M sodium acetate pH 4.6, incubated for 30 min at −20°C, centrifuged at 14.000 rpm 4°C for 30 min, washed with 70 % ethanol, and finally resuspended in 100 μl H_2_O. The integrity of the RNA was checked both in a 1.2% agarose gel and with RNA6000 Nano Assay using the Agilent 2100 Bioanalyzer (Agilent Technologies).

### RNA sequencing

Total RNA from *V. cholerae* co969 colonies grown at 37°C or 22°C was isolated as described above. RNA integrity was checked with RNA6000 Nano Assay using the Agilent 2100 Bioanalyzer (Agilent Technologies) and RNA samples were quantified using Qubit 2.0 Fluorometer (Life Technologies). To enrich the samples in mRNA, ribosomal RNA (rRNA) was removed from the samples using the Ribo-Zero rRNA removal kit (Illumina). cDNA preparation and sequencing reactions were conducted in the GeneCore Facility of the European Laboratory for Molecular Biology (EMBL, Heidelberg). Construction of mRNA seq libraries were conducted using the TruSeq Stranded Total RNA Library Prep Kit (Illumina) following the manufacturer’s recommendations. The samples were clustered on a flow cell and 50 cycle paired-end sequencing was performed on an Illumina HiSeq 2000 (Illumina).

For the data analysis, raw sequencing data generated from Illumina HiSeq2000 was converted into fastq files and de-multiplexed. Quality of the fastq files was checked with FastQC report and sequence reads were trimmed with Trimommatic^60^ to remove adapters, primers and reads with low qualities. The resulting sequence reads were aligned to the reference genome for *Vibrio cholerae* N16961 using Bowtie^61^ and the obtained .sam files were converted into .bam files with Samtools^62,63^. For each gene, sequence read account was calculated using custom Perl scripts. Differential gene expression analysis was performed with DESeq2/Bioconductor^64^. Principal component analysis was performed to examine sample separation between the two groups.

The number of reads per sample was: 5848844, 8628578 and 4899526 for samples 22 S1, 22 S2 and 22 S3; 27149959, 27241701 and 29623182 for samples 22 Sb1, 22 Sb2 and 22 Sb3; 7464802, 6855523, 7829998 for samples 37 S1, 37 S2 and 37 S3; and 8643032, 5549986 and 8006111 for samples 37 R1, 37 R2 and 37 R3. Mapping to *V. cholerae* N16961 genome resulted in the following numbers of aligned reads: 5656142, 8318183 and 4712069 for samples 22 S1, 22 S2 and 22 S3; 24917438, 24751935 and 26765757 for samples 22 Sb1, 22 Sb2 and 22 Sb3; 7144283, 6550527 and 7529534 for samples 37 S1, 37 S2 and 37 S3; and 8353239, 5375566 and 7727789 for samples 37 R1, 37 R2 and 37 R3, meaning a % of aligned reads between 90-97 %.

### Quantitative real-time PCR (qRT-PCR)

RNA was isolated from *V. cholera* colonies as described above. For cDNA synthesis, 4 μg of total RNA was incubated for 5 min at 65°C, to avoid formation of secondary structures, in the presence of dNTPs and random hexamer primers. Then, 200 U of Maxima RT (Thermo Scientific) and RNase inhibitor were added to the mix and further incubated 10 min at 25°C, 30 min at 50°C and finally 5 min at 85°C, for heat-inactivation of the enzyme. For RT-control reactions, to verify the absence of contaminant genomic DNA in the RNA samples, H_2_O was added instead of Maxima RT. Once the reactions were completed, approx. 5 U RNase A (Thermo Scientific) and 2.5 U RNase H (Thermo Scientific) were added to the reaction mixtures and incubated for 30 min at 37°C to remove the remaining DNA. The synthesized cDNA was purified using a QIAquick PCR purification kit (QIAGEN), and its concentration was determined spectrophotometrically in a Nano-drop Lite spectrophotometer (Thermo Scientific). Real time quantitative PCR (qRT-PCR) was performed using an iQ™5 Multicolor Real-Time PCR Detection System (BIO-RAD) with qPCRBIO SyGreen Mix fluorescein (PCR BIOSYSTEMS). Master mixes were prepared as recommended by the manufacturer, with qRT-PCR primers listed in SM Table 3. For normalization, either *gyrA* or *hfq* were used as internal standards.

### Transposon mutagenesis of *V. cholerae*

Transposon mutagenesis was performed by conjugation of *V. cholerae* co969 (recipient strain) with *E. coli* SM10 λ pir (pSC189)^65^ (donor strain), as described by Dörr et al.^66^. Transposon mutants were selected on LB plates (10 g/l NaCl) containing both streptomycin (200 μg/ml) and kanamycin (50 μg/ml). For the first screen, out of 11000 screened mutants, 459 were selected due to their rugose colony morphology at 22°C. To identify the genes interrupted by the transposon, the selected 459 mutants were pooled together and transposon insertion sequencing (Tn-seq) was performed as described by Chao et al.^67^. In brief, the chromosomal DNA of the mutants pooled together was extracted, DNA was sheared by sonication (Covaris sonicator E220), the overhanging ends were blunted using a blunting enzyme kit (NEB) and an A-tail was added with Taq DNA polymerase (NEB). Then, Illumina adaptors were ligated (with T4 DNA ligase, NEB) and the genomic DNA-transposon junctions were amplified (with Phusion^®^ High-Fidelity DNA polymerase, NEB). Sequencing was performed using an Illumina MiSeq benchtop sequencer (Illumina, San Diego, USA). Data analysis for determination of the TA insertion sites was conducted as described previously^67,68^. For visualization of transposon insertion profiles, Sanger Artemis Genome Browser and Annotation tool was used^69^.

To avoid selecting false positive candidates with a temperature-independent effect on biofilm formation, another transposon mutagenesis was performed in parallel, by conjugation of the constitutively smooth *V. cholerae* C6706 with *E. coli* SM10 λ pir (pSC189). Out of 63000 screened mutants, 618 formed rugose colonies at 37°C. Identification of the genes interrupted by the transposon was performed using the approach described above. Genes that appeared to be mutated in both transposon mutagenesis were disregarded during the selection of candidates for further analysis.

### Western blot

Samples for Western blotting were prepared as follows. 2 μl drops of *V. cholerae* co969:*bipA*-flag exponential cultures, started from a 1:100 dilution of an o/n culture, were plated on LB agar plates containing 10 g/l NaCl and incubated for an appropriate time at either 37°C or 22°C (under those conditions, the number of CFUs/colony was similar for both temperatures). For each sample, 2 colonies were collected, resuspended in buffer (20 mM Tris pH 7.5, 200 mM NaCl, 5 mM EDTA and protease inhibitor (cOmplete protease inhibitor cocktail tablets, Sigma Aldrich)), disrupted on a bead beater (Mini beadbeater, Biospec Products) (4°C, maximum speed for 1 min 15 sec) and the total protein concentration of the samples was determined by Bradford assay^70^ with Protein Assay Dye Reagent Concentrate (BIO-RAD). Samples were normalized by total protein content and run in a 10% SDS-PAGE gel for protein separation. Proteins were then transferred to an Immobilon-P transfer membrane (Millipore) using a Trans-Blot Turbo Transfer system (BIO-RAD). After blocking the membrane at room temperature for 2 h with TBS-T buffer (24.2 g Trizma, 87.6 g NaCl in 1 l H_2_O, pH 7.5, 0.1% (v/v) Tween-20) containing 5% (w/v) milk, the membrane was incubated at 4°C o/n with the monoclonal anti-flag M2 primary antibody (Sigma Aldrich) in TBS-T buffer containing 5% (w/v) milk, washed with TBS-T buffer, incubated at RT for 1 h with α-mouse secondary antibody in TBS-T buffer and again washed with TBS-T buffer. BipA-flag was detected by addition of SuperSignal^®^West Pico Chemiluminiscence substrate (Thermo Scientific) and detection of the signal using a LAS4000 (Fujifilm).

### Purification of BipA-His

*E. coli* BL21 cells carrying plasmid pET22b-*bipA*-His were grown at 37°C in LB media containing carbenicillin (100 μg/ml) and 0.4% glucose (w/v) until an OD_600_ of 0.4 was reached. Then, expression of the BipA-His fusion protein was induced by addition of 1mM IPTG (isopropyl-β-d-thiogalactosidase) and further incubation for 2 h at 37°C. Cells were then pelleted by centrifugation at 4°C 6000 rpm 30 min, resuspended in equilibration buffer (150 mM Tris-HCl pH 7.5, 200 mM NaCl) in the presence of protease inhibitors (cOmplete protease inhibitor cocktail tablets, Sigma Aldrich) and cell crude extracts were prepared by disruption in a pressure cell homogenizer FC600 (Julabo) followed by centrifugation at 4°C 15000 rpm 30 min. BipA-His was purified from the soluble fraction by affinity chromatography using Ni-NTA agarose (QIAGEN) previously equilibrated with equilibration buffer. After washing with washing buffer (150 mM Tris-HCl pH 7.5, 1 M NaCl), BipA-His was eluted with elution buffer (150 mM Tris-HCl pH 7.5, 200 mM NaCl, 500 mM imidazole) and then buffer exchanged to 100 mM Tris-HCl pH 7.5 + 100 mM NaCl by dialysis at 4°C o/n using a Spectra/Por^®^ molecular porous membrane tubing (6-8 kD) (Spectrumlabs.com). Purified proteins were either stored at 4°C, for immediate use, or at −80°C after addition of 10% glycerol. For circular dichroism experiments, one more buffer exchange was performed to 20 mM sodium phosphate buffer pH 7.5, 50 mM NaCl by dialysis at 4°C o/n using a Spectra/Por^®^ molecular porous membrane tubing (6-8 kD) (Spectrumlabs.com).

### Partial digestion with trypsin

Aliquots containing 20 μg of either purified BipA-His or BSA were pre-incubated for 2 h at 22°C or 37°C. Then, after incubation for 5 min at room temperature, samples were subjected to partial digestion by incubation with 1.5 μg trypsin for the indicated times: 0, 10 sec, 30 sec, 2 min and 5 min for BipA-His; or 0, 10, 30, 45, 60 and 90 min for BSA. The digested proteins were visualized in a 10% acrylamide gel stained with Coomassie Brilliant Blue R (Sigma Aldrich)

### Circular dichroism

2.5 μM of BipA-His in 20 mM sodium phosphate buffer pH 7.5, 50 mM NaCl were pre-incubated for 30 min at the appropriate temperature (37°C, 22°C or 15°C) or either pre-incubated for 30 min at 37°C and then sifted to 22°C for further 30 min. Then, circular dichroism was measured in a Jasco J-720 CD spectrometer (Jasco, Japan) (190-260 nm; 1 mm quartz cuvette, 3 μm scan, 5 averaged scans). Background spectra were always subtracted prior to analysis.

### Quantitative label-free proteomics

Samples used for quantitative label-free proteomics were prepared as follows. 2 μl drops of *V. cholerae* co969 or *V. cholerae* co969 Δ*bipA* o/d cultures, started from a 1:100 dilution of o/n cultures, were dispensed in LB agar plates containing 10 g/l NaCl and incubated the appropriate time at either 37°C or 22°C, so that the number of CFUs/colony was similar in both conditions. Three biological replicates, each of them being a pool of 4 independent colonies of *Vibrio cholerae* co969 or *V. cholerae* co969 Δ*bipA*, either grown at 37°C or 22°C (co969 37C, co969 22C, co969 Δ*bipA* 37C, co969 Δ*bipA* 22C) were analyzed. Total protein extracts were prepared by resuspending the collected pellets in 200 μl 20 mM Tris-HCl pH 7.5 + 200 mM NaCl, disrupting the cells in a bead beater (4°C, maximum speed for 1 min 15 sec) and collecting the supernatants. 20 μg of total protein extracts were run in a 12% pre-cast acrylamide gel at 100V for 20 min, and total protein bands were excised from the gel. Gel slices were digested as described previously^71^. Peptide mixtures were then separated on an EasyLC nano-HPLC (Proxeon Biosystems) coupled to an LTQ Orbitrap Elite mass spectrometer (Thermo Fisher Scientific) as described elsewhere^72^ with the following modifications: peptides were eluted with an 130-min segmented gradient of 5–33-90% HPLC solvent B (80% acetonitrile in 0.5% acetic acid). The acquired MS spectra were processed with the MaxQuant software package, version 1.5.2.8^73^ with the integrated Andromeda search engine^74^ as described previously^72^. Database searches were performed against a target-decoy *V. cholerae* complete database obtained from UniProt, containing 3,783 protein entries and 248 commonly observed contaminants. The label-free algorithm was enabled, as was the “match between runs” option^64^. Label-free quantification (LFQ) protein intensities from the MaxQuant data output were used for relative protein quantification. Downstream bioinformatic analysis (ANOVA and two-sample t-tests) was performed using the Perseus software package, version 1.5.0.15. P < 0.05 was considered to be statistically significant. Determination of statistically significant differences was only possible for proteins that appeared in all four conditions.

## Acknowledgements

We thank Laura Alvarez for the construction of plasmids pCVD442-Δ*hapR* and pCVD442-*hapR*c and Laura Alvarez and Alonso Serrano for assistance with RNA-seq data analysis We thank all the members of the Cava lab for helpful discussions. Proteomics were performed by Ana Velic, Nicolas Nalpas and Boris Mazek at the University of Tübingen (Germany). Research in the Cava lab is supported by The Swedish Research Council (VR), The Knut and Alice Wallenberg Foundation (KAW), The Laboratory of Molecular Infection Medicine Sweden (MIMS) and The Kempe Foundation. Research in the Waldor lab is supported by NIH grant RO1AI-042347 and HHMI. ARW was funded by grant T32 AI-132120. BS was supported by the Natural Sciences and Engineering Council of Canada (PGSD3-487259-2016). TdP was the recipient of an EMBO-short term fellowship (EMBO ASTF 1-2015).

## Author contributions

FC conceived the study. TdP, JB, BS, AW, VB, MW and FC designed the experiments. TdP performed the majority of experiments and analysis, except as noted. JB and VB performed and analysed RNAseq data, assisted by TdP. BS and AW helped with the phenotypical characterization of the *bipA* mutant. VB, MW and FC supervised the research. TdP and FC wrote the manuscript with support and editing from all authors.

## References

1 Islam, M. S. et al. Biofilm Acts as a Microenvironment for Plankton-Associated Vibrio cholerae in the Aquatic Environment of Bangladesh. Microbiology and Immunology 51, 369–379, doi:10.1111/j.1348-0421.2007.tb03924.x (2007).

2 Alam, M. et al. Viable but nonculturable Vibrio cholerae O1 in biofilms in the aquatic environment and their role in cholera transmission. Proceedings of the National Academy of Sciences of the United States of America 104, 17801–17806, doi:10.1073/pnas.0705599104 (2007).

3 Lutz, C., Erken, M., Noorian, P., Sun, S. & McDougald, D. Environmental reservoirs and mechanisms of persistence of Vibrio cholerae. Front Microbiol 4, 375, doi:10.3389/fmicb.2013.00375 (2013).

4 Faruque, S. M. et al. Transmissibility of cholera: in vivo-formed biofilms and their relationship to infectivity and persistence in the environment. Proceedings of the National Academy of Sciences of the United States of America 103, 6350–6355, doi:10.1073/pnas.0601277103 (2006).

5 Silva, A. J. & Benitez, J. A. Vibrio cholerae Biofilms and Cholera Pathogenesis. PLoS Negl Trop Dis 10, e0004330, doi:10.1371/journal.pntd.0004330 (2016).

6 Tamayo, R., Patimalla, B. & Camilli, A. Growth in a biofilm induces a hyperinfectious phenotype in Vibrio cholerae. Infect Immun 78, 3560–3569, doi:10.1128/IAI.00048-10 (2010).

7 Fong, J. C. N., Syed, K. A., Klose, K. E. & Yildiz, F. H. Role of Vibrio polysaccharide (vps) genes in VPS production, biofilm formation and Vibrio cholerae pathogenesis. Microbiology 156, 2757–2769, doi:10.1099/mic.0.040196-0 (2010).

8 Yildiz, F. H. & Schoolnik, G. K. Vibrio cholerae O1 El Tor: identification of a gene cluster required for the rugose colony type, exopolysaccharide production, chlorine resistance, and biofilm formation. Proc Natl Acad Sci U S A 96, 4028–4033, doi:10.1073/pnas.96.7.4028 (1999).

9 Fong, J. C. N., Karplus, K., Schoolnik, G. K. & Yildiz, F. H. Identification and characterization of RbmA, a novel protein required for the development of rugose colony morphology and biofilm structure in Vibrio cholerae. Journal of bacteriology 188, 1049–1059, doi:10.1128/JB.188.3.1049-1059.2006 (2006).

10 Fong, J. C. & Yildiz, F. H. The rbmBCDEF gene cluster modulates development of rugose colony morphology and biofilm formation in Vibrio cholerae. J Bacteriol 189, 2319–2330, doi:10.1128/JB.01569-06 (2007).

11 Reichhardt, C., Fong, J. C., Yildiz, F. & Cegelski, L. Characterization of the Vibrio cholerae extracellular matrix: a top-down solid-state NMR approach. Biochim Biophys Acta 1848, 378–383, doi:10.1016/j.bbamem.2014.05.030 (2015).

12 Berk, V. et al. Molecular architecture and assembly principles of Vibrio cholerae biofilms. Science 337, 236–239, doi:10.1126/science.1222981 (2012).

13 Seper, A. et al. Extracellular nucleases and extracellular DNA play important roles in Vibrio cholerae biofilm formation. Mol Microbiol 82, 1015–1037, doi:10.1111/j.1365-2958.2011.07867.x (2011).

14 Teschler, J. K. et al. Living in the matrix: assembly and control of Vibrio cholerae biofilms. Nat Rev Microbiol 13, 255–268, doi:10.1038/nrmicro3433 (2015).

15 Conner, J. G., Teschler, J. K., Jones, C. J. & Yildiz, F. H. Staying Alive: Vibrio cholerae’s Cycle of Environmental Survival, Transmission, and Dissemination. Microbiol Spectr 4, doi:10.1128/microbiolspec.VMBF-0015-2015 (2016).

16 Beyhan, S., Bilecen, K., Salama, S. R., Casper-Lindley, C. & Yildiz, F. H. Regulation of rugosity and biofilm formation in Vibrio cholerae: comparison of VpsT and VpsR regulons and epistasis analysis of vpsT, vpsR, and hapR. J Bacteriol 189, 388–402, doi:10.1128/JB.00981-06 (2007).

17 Zamorano-Sanchez, D., Fong, J. C., Kilic, S., Erill, I. & Yildiz, F. H. Identification and characterization of VpsR and VpsT binding sites in Vibrio cholerae. J Bacteriol 197, 1221–1235, doi:10.1128/JB.02439-14 (2015).

18 Waters, C. M., Lu, W., Rabinowitz, J. D. & Bassler, B. L. Quorum sensing controls biofilm formation in Vibrio cholerae through modulation of cyclic di-GMP levels and repression of vpsT. Journal of bacteriology 190, 2527–2536, doi:10.1128/JB.01756-07 (2008).

19 Zhu, J. et al. Quorum-sensing regulators control virulence gene expression in Vibrio cholerae. Proc Natl Acad Sci U S A 99, 3129–3134, doi:10.1073/pnas.052694299 (2002).

20 Kovacikova, G. & Skorupski, K. Regulation of virulence gene expression in Vibrio cholerae by quorum sensing: HapR functions at the aphA promoter. Molecular Microbiology 46, 1135–1147, doi:10.1046/j.1365-2958.2002.03229.x (2002).

21 Zheng, J., Shin, O. S., Cameron, D. E. & Mekalanos, J. J. Quorum sensing and a global regulator TsrA control expression of type VI secretion and virulence in Vibrio cholerae. Proceedings of the National Academy of Sciences of the United States of America 107, 21128–21133, doi:10.1073/pnas.1014998107 (2010).

22 Hammer, B. K. & Bassler, B. L. Distinct sensory pathways in Vibrio cholerae El Tor and classical biotypes modulate cyclic dimeric GMP levels to control biofilm formation. J Bacteriol 191, 169–177, doi:10.1128/JB.01307-08 (2009).

23 Hammer, B. K. & Bassler, B. L. Quorum sensing controls biofilm formation in Vibrio cholerae. Mol Microbiol 50, 101–104, doi:10.1046/j.1365-2958.2003.03688.x (2003).

24 Zhu, J. & Mekalanos, J. J. Quorum sensing-dependent biofilms enhance colonization in Vibrio cholerae. Dev Cell 5, 647–656, doi:10.1016/s1534-5807(03)00295-8 (2003).

25 Tsou, A. M., Liu, Z., Cai, T. & Zhu, J. The VarS/VarA two-component system modulates the activity of the Vibrio cholerae quorum-sensing transcriptional regulator HapR. Microbiology 157, 1620–1628, doi:10.1099/mic.0.046235-0 (2011).

26 Lenz, D. H. & Bassler, B. L. The small nucleoid protein Fis is involved in Vibrio cholerae quorum sensing. Mol Microbiol 63, 859–871, doi:10.1111/j.1365-2958.2006.05545.x (2007).

27 Shikuma, N. J. et al. Overexpression of VpsS, a hybrid sensor kinase, enhances biofilm formation in Vibrio cholerae. J Bacteriol 191, 5147–5158, doi:10.1128/JB.00401-09 (2009).

28 Liang, W., Pascual-Montano, A., Silva, A. J. & Benitez, J. A. The cyclic AMP receptor protein modulates quorum sensing, motility and multiple genes that affect intestinal colonization in Vibrio cholerae. Microbiology 153, 2964–2975, doi:https://doi.org/10.1099/mic.0.2007/006668-0 (2007).

29 Liu, Z., Hsiao, A., Joelsson, A. & Zhu, J. The transcriptional regulator VqmA increases expression of the quorum-sensing activator HapR in Vibrio cholerae. J Bacteriol 188, 2446–2453, doi:10.1128/JB.188.7.2446-2453.2006 (2006).

30 Yildiz, F. H., Liu, X. S., Heydorn, A. & Schoolnik, G. K. Molecular analysis of rugosity in a Vibrio cholerae O1 El Tor phase variant. Mol Microbiol 53, 497–515, doi:10.1111/j.1365-2958.2004.04154.x (2004).

31 Joelsson, A., Liu, Z. & Zhu, J. Genetic and phenotypic diversity of quorum-sensing systems in clinical and environmental isolates of Vibrio cholerae. Infect Immun 74, 1141–1147, doi:10.1128/IAI.74.2.1141-1147.2006 (2006).

32 Chowdhury, G. et al. Rugose atypical Vibrio cholerae O1 El Tor responsible for 2009 cholera outbreak in India. J Med Microbiol 65, 1130–1136, doi:10.1099/jmm.0.000344 (2016).

33 Katzianer, D. S., Wang, H., Carey, R. M. & Zhu, J. “Quorum Non-Sensing”: Social Cheating and Deception in Vibrio cholerae. Appl Environ Microbiol 81, 3856–3862, doi:10.1128/AEM.00586-15 (2015).

34 Parsot, C. & Mekalanos, J. J. Expression of ToxR, the transcriptional activator of the virulence factors in Vibrio cholerae, is modulated by the heat shock response. Proceedings of the National Academy of Sciences of the United States of America 87, 9898–9902, doi:10.1073/pnas.87.24.9898 (1990).

35 Weber, G. G., Kortmann, J., Narberhaus, F. & Klose, K. E. RNA thermometer controls temperature-dependent virulence factor expression in Vibrio cholerae. Proc Natl Acad Sci U S A 111, 14241–14246, doi:10.1073/pnas.1411570111 (2014).

36 Margus, T., Remm, M. & Tenson, T. Phylogenetic distribution of translational GTPases in bacteria. BMC Genomics 8, 15, doi:10.1186/1471-2164-8-15 (2007).

37 Leipe, D. D., Wolf, Y. I., Koonin, E. V. & Aravind, L. Classification and evolution of P-loop GTPases and related ATPases. J Mol Biol 317, 41–72, doi:10.1006/jmbi.2001.5378 (2002).

38 Grant, A. J. et al. Co-ordination of pathogenicity island expression by the BipA GTPase in enteropathogenic Escherichia coli (EPEC). Mol Microbiol 48, 507–521, doi:10.1046/j.1365-2958.2003.t01-1-03447.x (2003).

39 Farris, M., Grant, A., Richardson, T. B. & O’Connor, C. D. BipA: a tyrosine-phosphorylated GTPase that mediates interactions between enteropathogenic Escherichia coli (EPEC) and epithelial cells. Molecular Microbiology 28, 265–279, doi:10.1046/j.1365-2958.1998.00793.x (1998).

40 deLivron, M. A. & Robinson, V. L. Salmonella enterica serovar Typhimurium BipA exhibits two distinct ribosome binding modes. J Bacteriol 190, 5944–5952, doi:10.1128/JB.00763-08 (2008).

41 Neidig, A. et al. TypA is involved in virulence, antimicrobial resistance and biofilm formation in Pseudomonas aeruginosa. BMC Microbiol 13, 77–77, doi:10.1186/1471-2180-13-77 (2013).

42 Kiss, E., Huguet, T., Poinsot, V. & Batut, J. The typA Gene is Required for Stress Adaptation as Well as for Symbiosis of Sinorhizobium meliloti 1021 with Certain Medicago truncatula Lines. Molecular Plant-Microbe Interactions® 17, 235–244, doi:10.1094/MPMI.2004.17.3.235 (2004).

43 Beckering, C. L., Steil, L., Weber, M. H., Volker, U. & Marahiel, M. A. Genomewide transcriptional analysis of the cold shock response in Bacillus subtilis. J Bacteriol 184, 6395–6402, doi:10.1128/jb.184.22.6395-6402.2002 (2002).

44 Pfennig, P. & Flower, A. BipA is required for growth of Escherichia coli K12 at low temperature. Molecular Genetics and Genomics 266, 313–317, doi:10.1007/s004380100559 (2001).

45 Hiramatsu, Y. et al. BipA Is Associated with Preventing Autoagglutination and Promoting Biofilm Formation in Bordetella holmesii. PLoS One 11, e0159999–e0159999, doi:10.1371/journal.pone.0159999 (2016).

46 Overhage, J., Lewenza, S., Marr, A. K. & Hancock, R. E. W. Identification of genes involved in swarming motility using a Pseudomonas aeruginosa PAO1 mini-Tn5-lux mutant library. Journal of bacteriology 189, 2164–2169, doi:10.1128/JB.01623-06 (2007).

47 Fan, H., Hahm, J., Diggs, S., Perry, J. J. & Blaha, G. Structural and Functional Analysis of BipA, a Regulator of Virulence in Enteropathogenic Escherichia coli. J Biol Chem 290, 20856–20864, doi:10.1074/jbc.M115.659136 (2015).

48 Rogers, A. et al. The LonA Protease Regulates Biofilm Formation, Motility, Virulence, and the Type VI Secretion System in Vibrio cholerae. J Bacteriol 198, 973–985, doi:10.1128/JB.00741-15 (2016).

49 Gibbs, M. R. & Fredrick, K. Roles of elusive translational GTPases come to light and inform on the process of ribosome biogenesis in bacteria. Mol Microbiol 107, 445–454, doi:10.1111/mmi.13895 (2018).

50 Starosta, A. L., Lassak, J., Jung, K. & Wilson, D. N. The bacterial translation stress response. FEMS Microbiol Rev 38, 1172–1201, doi:10.1111/1574-6976.12083 (2014).

51 Krishnan, K. & Flower, A. M. Suppression of DeltabipA phenotypes in Escherichia coli by abolishment of pseudouridylation at specific sites on the 23S rRNA. Journal of bacteriology 190, 7675–7683, doi:10.1128/JB.00835-08 (2008).

52 Choudhury, P. & Flower, A. M. Efficient assembly of ribosomes is inhibited by deletion of bipA in Escherichia coli. Journal of bacteriology 197, 1819–1827, doi:10.1128/JB.00023-15 (2015).

53 Watnick, P. I. & Kolter, R. Steps in the development of a Vibrio cholerae El Tor biofilm. Mol Microbiol 34, 586–595, doi:10.1046/j.1365-2958.1999.01624.x (1999).

54 Townsley, L. & Yildiz, F. H. Temperature affects c-di-GMP signalling and biofilm formation in Vibrio cholerae. Environ Microbiol 17, 4290–4305, doi:10.1111/1462-2920.12799 (2015).

55 Townsley, L., Sison Mangus, M. P., Mehic, S. & Yildiz, F. H. Response of Vibrio cholerae to Low-Temperature Shifts: CspV Regulation of Type VI Secretion, Biofilm Formation, and Association with Zooplankton. Appl Environ Microbiol 82, 4441–4452, doi:10.1128/AEM.00807-16 (2016).

56 Qi, S. Y. et al. Salmonella typhimurium responses to a bactericidal protein from human neutrophils. Mol Microbiol 17, 523–531, doi:10.1111/j.1365-2958.1995.mmi_17030523.x (1995).

57 Barker, H. C., Kinsella, N., Jaspe, A., Friedrich, T. & O’Connor, C. D. Formate protects stationary-phase Escherichia coli and Salmonella cells from killing by a cationic antimicrobial peptide. Mol Microbiol 35, 1518–1529, doi:10.1046/j.1365-2958.2000.01820.x (2000).

58 Mathur, J. & Waldor, M. K. The Vibrio cholerae ToxR-regulated porin OmpU confers resistance to antimicrobial peptides. Infect Immun 72, 3577–3583, doi:10.1128/IAI.72.6.3577-3583.2004 (2004).

59 Schneider, C. A., Rasband, W. S. & Eliceiri, K. W. NIH Image to ImageJ: 25 years of image analysis. Nat Methods 9, 671–675, doi:10.1038/nmeth.2089 (2012).

60 Bolger, A. M., Lohse, M. & Usadel, B. Trimmomatic: a flexible trimmer for Illumina sequence data. Bioinformatics 30, 2114–2120, doi:10.1093/bioinformatics/btu170 (2014).

61 Langmead, B. & Salzberg, S. L. Fast gapped-read alignment with Bowtie 2. Nat Methods 9, 357–359, doi:10.1038/nmeth.1923 (2012).

62 Li, H. et al. The Sequence Alignment/Map format and SAMtools. Bioinformatics (Oxford, England) 25, 2078–2079, doi:10.1093/bioinformatics/btp352 (2009).

63 Li, H. A statistical framework for SNP calling, mutation discovery, association mapping and population genetical parameter estimation from sequencing data. Bioinformatics 27, 2987–2993, doi:10.1093/bioinformatics/btr509 (2011).

64 Luber, C. A. et al. Quantitative proteomics reveals subset-specific viral recognition in dendritic cells. Immunity 32, 279–289, doi:10.1016/j.immuni.2010.01.013 (2010).

65 Chiang, S. L. & Rubin, E. J. Construction of a mariner-based transposon for epitope-tagging and genomic targeting. Gene 296, 179–185, doi:10.1016/s0378-1119(02)00856-9 (2002).

66 Dorr, T. et al. A Transposon Screen Identifies Genetic Determinants of Vibrio cholerae Resistance to High-Molecular-Weight Antibiotics. Antimicrob Agents Chemother 60, 4757–4763, doi:10.1128/AAC.00576-16 (2016).

67 Chao, M. C. et al. High-resolution definition of the Vibrio cholerae essential gene set with hidden Markov model-based analyses of transposon-insertion sequencing data. Nucleic Acids Res 41, 9033–9048, doi:10.1093/nar/gkt654 (2013).

68 Pritchard, J. R. et al. ARTIST: high-resolution genome-wide assessment of fitness using transposon-insertion sequencing. PLoS Genet 10, e1004782–e1004782, doi:10.1371/journal.pgen.1004782 (2014).

69 Rutherford, K. et al. Artemis: sequence visualization and annotation. Bioinformatics 16, 944–945, doi:10.1093/bioinformatics/16.10.944 (2000).

70 Bradford, M. M. A rapid and sensitive method for the quantitation of microgram quantities of protein utilizing the principle of protein-dye binding. Anal Biochem 72, 248–254, doi:10.1006/abio.1976.9999 (1976).

71 Burian, M. et al. Quantitative proteomics of the human skin secretome reveal a reduction in immune defense mediators in ectodermal dysplasia patients. J Invest Dermatol 135, 759–767, doi:10.1038/jid.2014.462 (2015).

72 Carpy, A. et al. Absolute proteome and phosphoproteome dynamics during the cell cycle of Schizosaccharomyces pombe (Fission Yeast). Mol Cell Proteomics 13, 1925–1936, doi:10.1074/mcp.M113.035824 (2014).

73 Cox, J. & Mann, M. MaxQuant enables high peptide identification rates, individualized p.p.b.-range mass accuracies and proteome-wide protein quantification. Nature biotechnology 26, 1367–1372, doi:10.1038/nbt.1511 (2008).

74 Cox, J. et al. Andromeda: a peptide search engine integrated into the MaxQuant environment. Journal of proteome research 10, 1794–1805, doi:10.1021/pr101065j (2011).

